# Hi-Fi fMRI: High-resolution, fast-sampled and sub-second whole-brain functional MRI at 3T in humans

**DOI:** 10.1101/2023.05.13.540663

**Authors:** Benedetta Franceschiello, Simone Rumac, Mauro Leidi, Tom Hilbert, Matthias Nau, Martyna Dziadosz, Giulio Degano, Christopher W. Roy, Anna Gaglianese, Giovanni Petri, Jérôme Yerly, Matthias Stuber, Tobias Kober, Ruud B. van Heeswijk, Jessica A.M. Bastiaansen, João Jorge, Micah M. Murray, Eleonora Fornari

**Author notes:** Shared senior authorship (Micah M. Murray, Eleonora Fornari).

## Abstract

Functional magnetic resonance imaging (fMRI) is a methodological cornerstone of neuroscience. Most studies measure blood-oxygen-level-dependent (BOLD) signal using echo-planar imaging (EPI), Cartesian sampling, and image reconstruction with a one-to-one correspondence between the number of acquired volumes and reconstructed images. However, EPI schemes are subject to trade-offs between spatial and temporal resolutions. We make strides in overcoming these limitations by measuring BOLD with a gradient recalled echo (GRE) with a 3D radial-spiral phyllotaxis trajectory at a high sampling rate (28.49ms) on standard 3T field strength. The framework enables the reconstruction of 3D signal time courses with whole-brain coverage at simultaneously higher nominal spatial (1mm^3^) and temporal (up to 250ms) resolutions, as compared to optimized EPI schemes. Additionally, we apply motion correction directly to the k-space raw data, enabling flexible motion-corrected reconstructions; the desired temporal resolution to observe hemodynamic responses can be chosen after scanning. By showing activation in the calcarine sulcus of 20 participants completing an ON-OFF visual paradigm, we demonstrate the reliability of our method for applications in cognitive neuroscience research.

## 1 Introduction

The discovery of the blood-oxygen-level-dependent (BOLD) signal in the 1990s (1; 2; 3; 4) made it possible to non-invasively measure functional brain activity with Magnetic Resonance Imaging (MRI), which has since been a major driver of advances in human cognitive neuroscience. Subsequently, extensive research has been devoted towards developing new methods and improvements for fMRI acquisition with the goal of increasing functional sensitivity, as well as temporal and spatial specificity. However, like any other technique, BOLD fMRI has its limitations, and users must often compromise between temporal versus spatial resolutions alongside the extent of brain coverage (i.e., field of view; FoV) (5).

The introduction of echo-planar imaging (EPI) (6; 7; 8) made it possible to acquire images across the entire brain with reasonable compromises in terms of speed, spatial resolution, and sensitivity to artifacts (3; 4). EPI (8) proved early on to be an excellent option for BOLD fMRI in humans (3; 4), and remains the most widely adopted approach for fMRI. Its 2-dimensional (2D) form, where one 2D slice is acquired for each radio-frequency (RF) excitation, has been improved with in-plane acceleration techniques by subsampling the frequency domain (k-space) (9), while leveraging differences in spatial information provided by multiple receiver elements of RF arrays (coils). Additionally, multi-slice acquisition techniques have been introduced to capture multiple slices for each excitation (10) and non-Cartesian trajectories in the 2D plane have also been developed (11) to improve the trade-off between spatial and temporal resolutions. In parallel, extensions of this technique to 3D k-space have also been extensively explored, namely with segmented acquisitions where one plane of 3D k-space is acquired per RF excitation, allowing under-sampling both in-plane and through-plane (12; 13). In general, the acquisition of one plane per excitation has been found to be a favorable choice for BOLD fMRI, because at typical *B*_0_ strengths (1.5–7T) the sensitivity of this contrast peaks at relatively long echo-time (TE) values (i.e., tens of milliseconds), which fits well with the relatively long planar sampling scheme. EPI can be affected by important artifacts such as ghosting and spatial distortions (14), and the requirement of multiple RF excitations to cover the volume of interest introduces important constraints in sampling speed. For example, baseline EPI schemes take 2–3s to acquire 3mm isotropic images of the whole brain at 3T magnetic field strength (15).

To break new ground in terms of acquisition speed and flexibility, 3D acquisitions have been pushed to highly accelerated, “non-planar” forms where the full 3D k-space can be sampled in a single RF excitation – also called echo volumar imaging (EVI) (7; 16). The acceleration relies on differences in spatial sensitivity across multiple receive elements of RF arrays. In extremis, the acquisitions can be performed with no gradient encoding whatsoever (in other words, sampling only the central part of the k-space) and rely purely on the differentiation allowed by the different array coils (17; 18; 19), in analogy with another imaging modality, magnetoencephalography (MEG). This concept has been termed magnetic resonance encephalography (MREG) by some groups (19), as well as inverse imaging (InI) by others (20). To partially mitigate their rather limited spatial specificity, the developments typically led to hybrid solutions with some degree of gradient-based encoding, using optimized non-Cartesian 3D trajectories, which can still capture the fastest physiological fluctuations of interest (18; 19; 21) (Altogether, the focus of these approaches nonetheless remains on temporal over spatial resolution).

While the above approaches can strongly leverage information redundancies in the spatial domain, functional hemodynamic fluctuations are likewise not randomly distributed over time, which can be exploited to further advance fMRI acquisition techniques. This idea has been leveraged in approaches such as hybrid radial-cartesian 3D EPI, where each k-space plane sampling is defined by rotating the previous one by the golden-angle around a fixed axis. In this way, 3D volumes can then be flexibly reconstructed using a variable number of consecutively-acquired planes, in a trade-off between temporal resolution and image quality (22). Others have retained the use of Cartesian grids, but explored methods such as k-t or compressed sensing to leverage sampling redundancies over k-space and time, with specialized techniques for reconstruction of the full spatiotemporal data. One such example is k-t FASTER, which employs low-rank matrix completion to leverage these redundancies (23). Finally, another important source of temporal dependencies can be introduced by the experimental paradigm itself, when repeated trials of the same stimulus/response are elicited during the acquisition. If appropriately timed, different fMRI samples from different response trials can be re-ordered with respect to the trial onset to obtain a combined response waveform at higher temporal resolution; this concept has been applied by numerous fMRI studies (24; 25; 26), but typically using conventional 2D EPI samples, which do not leverage other signal redundancies across time. Other technical advances have been shown to measure more fine-grained brain dynamics based on the local influence of electric activity on the magnetic field (27), which currently rely on ultra-high field strength (9.4T), are prone to motion artifacts, and were tested only in anesthetized animals on well-studied neurobiological systems (i.e., visual sensory cortices or whisker fields in rodents) (27; 28). Such approaches have been referred to as ultra-fast fMRI and can be powerfully combined with modeling or optogenetic techniques (29; 30). Nevertheless, scaling such techniques to study the awake human brain to measure BOLD activity remains extremely challenging and the robustness of the methods is currently debated (31).

Here, we introduce Hi-Fi fMRI, a framework operating on a standard clinical MRI system that achieves a 3D k-space acquisition sampling rate of 28.49ms, whole-brain coverage, and 1mm isotropic nominal spatial resolution. Hi-Fi fMRI is based on an efficient pseudo-randomized, non-Cartesian 3D sampling technique inspired by cardiovascular imaging (32), where spatiotemporal redundancies are commonly leveraged to obtain high-quality depictions of the movement of the heart or the dynamic distribution of injected contrast agent (33), over short scanning periods. Using concepts of compressed sensing for reconstruction, this technique allows flexible combination (or binning) of k-space readouts as a trade-off between temporal resolution and image quality. In this way, temporal resolution is set at reconstruction (that is, after scanning), hence allowing observation of the hemodynamic response at different temporal resolutions. Moreover, the percent signal change curve can be studied at each such different temporal resolution. To pursue high-quality reconstructions with both high spatial and temporal resolutions, we apply this technique in a task-based fMRI design with repeated stimulation trials (in direct analogy to repeated heartbeat cycles), allowing for flexible post-hoc combination of the pseudo-randomized 3D k-space samples. The Hi-Fi fMRI framework represents a first proof of concept. It aims to provide a pathway to overcome limitations of the quasi-standard and widely used EPI-based BOLD imaging technique, and studying human brain dynamics from a different angle.

## 2 Results

### 2.1 High spatial and temporal resolutions in BOLD contrast maps

Twenty healthy volunteers underwent a single MRI session in a 3T clinical scanner (MAGNETOM Prisma *^fit^*, Siemens Healthcare, Erlangen, Germany) as part of an established blocked-design flickering checkerboard visual stimulation experiment (34), fig. 1, top row left. Data were acquired using an uninterrupted gradient recalled echo (GRE) sequence with a 3D radial-spiral phyllotaxis sampling trajectory (35) (fig. 1, top row center). Each k-space readout was acquired with a TR of 28.49ms and a voxel size of 1mm^3^ (nominal spatial resolution). The versatility of the framework lies in the fact that single readouts acquired in the image sampling space (k-space) can be grouped together (i.e., “binned”) flexibly and retrospectively.

**Figure 1.**
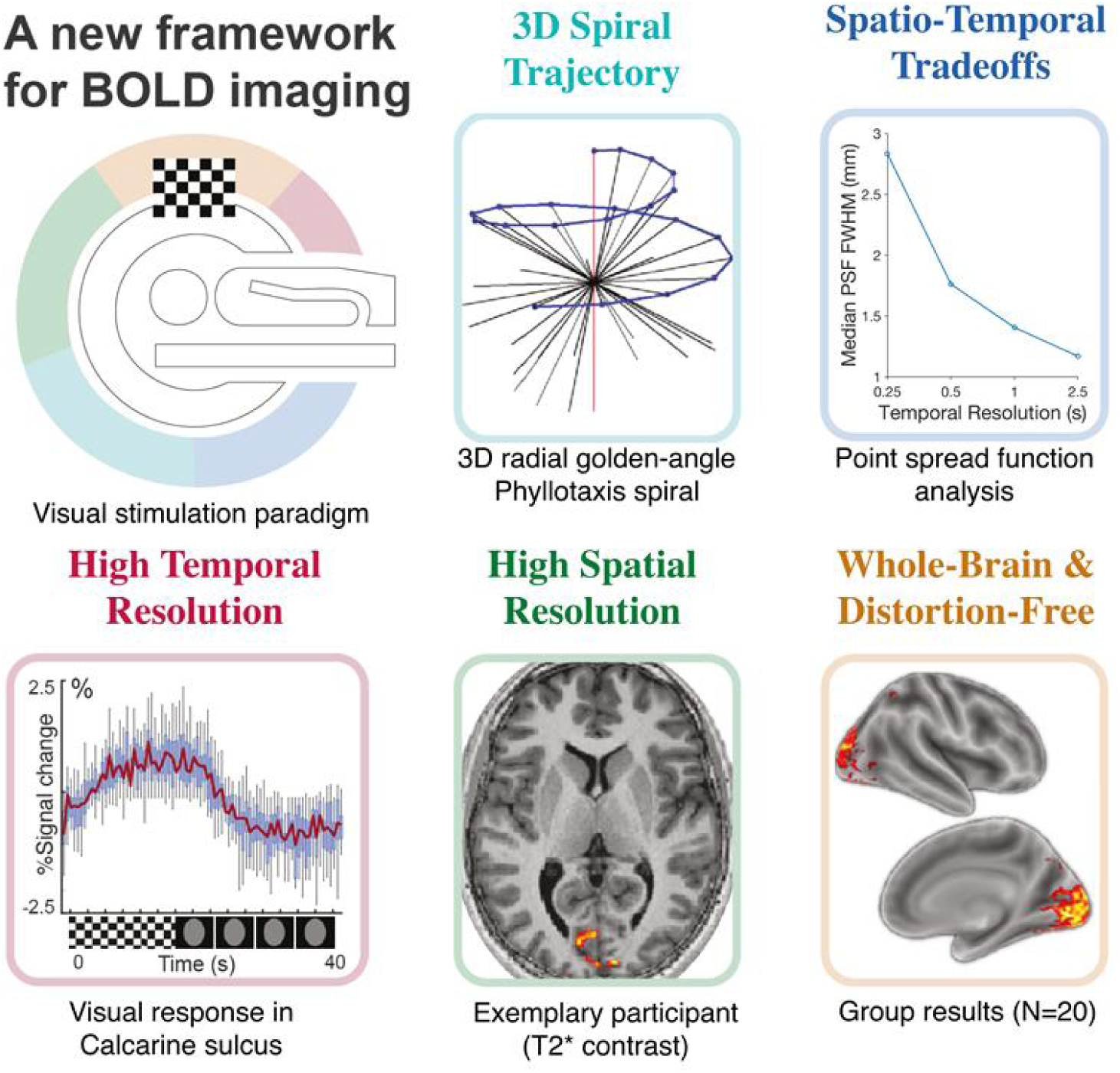
Graphical summary. Data were acquired using an uninterrupted Gradient Recalled Echo (GRE) sequence with a 3D radial spiral phyllotaxis sampling trajectory (top center), and a single readout TR of 28.49ms, while participants underwent a block design flickering checkerboard visual stimulation, alternating 15s ON active stimulation and 25s OFF resting periods. Different temporal image reconstructions were performed to track the evolution of the percent signal change across the 40-second trial interval at different temporal resolutions: 2.5s, 1s, 500ms, 250ms. The Point Spread Function (PSF) was estimated using AFNI from the spatial autocorrelation of the residuals of a General Linear Model (GLM) fitted to the single voxel time courses. Increasing temporal resolution was associated with higher FWHM values, increasing from 1.17 mm at 2.5 s to 2.83 mm at 250 ms. The bottom central figure shows the highly localized activation we observe at the single-participant level (calcarine, occipital cortex, at 1mm^3^ nominal spatial resolution). Group-level activation (bottom right) was obtained via a paired t-test (*p <* 0.001, extent threshold of 100 contiguous voxels).

This capability enables computation of percent signal change associated with an experimental manipulation at a finer temporal scale retrospectively, i.e. during reconstruction and thus after acquisition. Furthermore, T^∗^ BOLD contrast maps can be computed at the single-participant level as the mathematical difference I*ON* − *OFF*I, or as a one-tailed paired-t-test, see figure 7, p < 0.001, extended threshold of 100 contiguous voxels. Here, *ON* refers to the volume obtained through reconstruction of all readouts acquired during the stimulus-evoked hemodynamic response function peak (assumed to reach a plateau 5 seconds after stimulus onset and to last for 10 seconds in correspondence with our stimulation), while *OFF* refers to the volume obtained unifying all readouts acquired during the resting phase of the response (assumed to be 25-35 seconds post-stimulus) (36).

The obtained contrast maps replicate their classical counterpart such as those estimated via a canonical General Linear Model (GLM) analysis and EPI acquisition (37; 38). We generated 1 mm^3^ activation contrast maps with a whole-brain coverage for each participant individually, consistently showing a highly localized activation at the calcarine sulcus along the occipital lobe as expected with the task (see fig. 1, bottom row, center for an exemplar, individual participant). Voxel-wise group-level analysis (fig. 1, bottom row right) confirmed the single-participant results. In addition, we performed a canonical statistical analysis on one subject (single-subject level), showing that it is possible to inspect thresholded maps, as conventionally done in the field to investigate new activation areas or paradigms (supplementary materials, fig. 7).

### 2.2 Percent BOLD-signal change in the visual cortex with a temporal resolution up to 250ms

Using the same data, we reconstructed average trial time course images at different temporal resolutions, (fig. 1, top row right - Point Spread Function (PSF) analysis), grouping the signal across all trials at 250ms, 500ms, 1s, and 2.5s temporal resolutions, respectively.

Percent signal change was reliably extracted at different temporal resolutions within the calcarine sulcus, which includes primary and secondary visual areas. The measured percent signal change (1%, fig. 2 (a), left) indicates a consistent hemodynamic response in the calcarine sulcus across different temporal resolutions. By contrast, no reliable activation was observed in any of the other control ROIs, including the superior temporal lobe and precuneus (<0.05%, fig. 2, b). These results replicate the well-known response to visual stimulation in and around the calcarine sulcus (37; 38; 39; 40). Despite the increased number of bins (and conversely the decreased number of readouts per bin), the recorded percent signal change remained stable, with estimated PSF below 3mm, see (fig. 1, Spatio-Temporal Tradeoffs).

**Figure 2.**
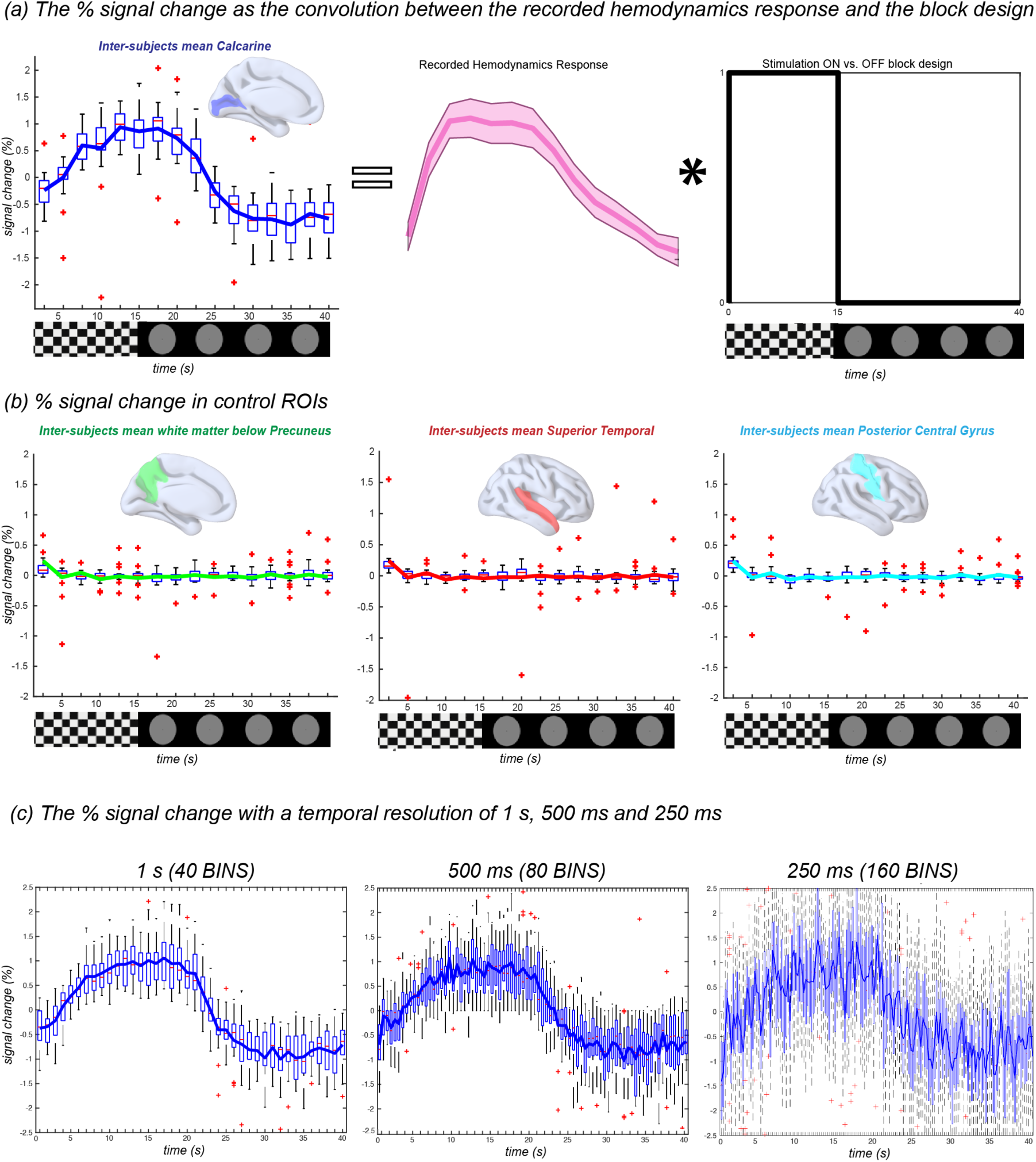
Percent BOLD-signal change in the visual cortex with a temporal resolution up to 250ms. In the top row, we show the percent signal changes in the calcarine sulcus, estimated from 4D reconstruction at a temporal resolution of 2.5s (16 bins). We retrieve the canonical hemodynamic response (center) as extracted by deconvolving the block design (right). The middle row reports the extraction of the percent signal change in different ROIs (Precuneus - green, Temporal Lobe - red, Posterior Central Gyrus - cyan). As expected, activity within these ROIs remains below significance threshold (<0.05%) as these regions are not a priori expected to respond to visual stimuli. The bottom row contains the extracted percent signal change at different temporal resolutions (averaged across trials at a temporal resolution of 1s, 500ms, or 250ms in a corresponding number of 40, 80, or 160 bins). Despite the increased temporal resolution, the recorded percent signal change remains stable, with estimated PSF below 3mm, see (fig. 1, Spatio-Temporal Tradeoffs).. Solid colored lines represent the averaged signal across voxels in the respective ROIs for each time point across subjects; each value on the x-axis (time with respect to stimulation) is therefore described by its mean, the corresponding box plot across subjects and the outliers (red crosses).

The PSF was estimated using Analysis of Functional NeuroImages (AFNI) from the spatial autocorrelation of the residuals of a General Linear Model (GLM) fitted to the voxel time courses at different temporal resolutions. Increasing temporal resolution was associated with higher Full Width at Half Maximum (FWHM) values, highlighting the effects of under-sampling on the effective spatial resolution of the reconstructed images. Estimated median values of FWHM were 1.17 mm, 1.41mm, 1.76mm and 2.83mm at 2.5s, 1s, 500ms and 250ms respectively (fig. 1, top row right - Point Spread Function (PSF) analysis and red histogram in fig. 3, section C). The under-sampling rate is proportional to the temporal resolution and inversely proportional to the number of trials. Consequently, this framework mitigates the traditional trade-off between temporal and spatial resolution by replacing it with a trade-off between the number of repeated trials and spatial resolution.

**Figure 3.**
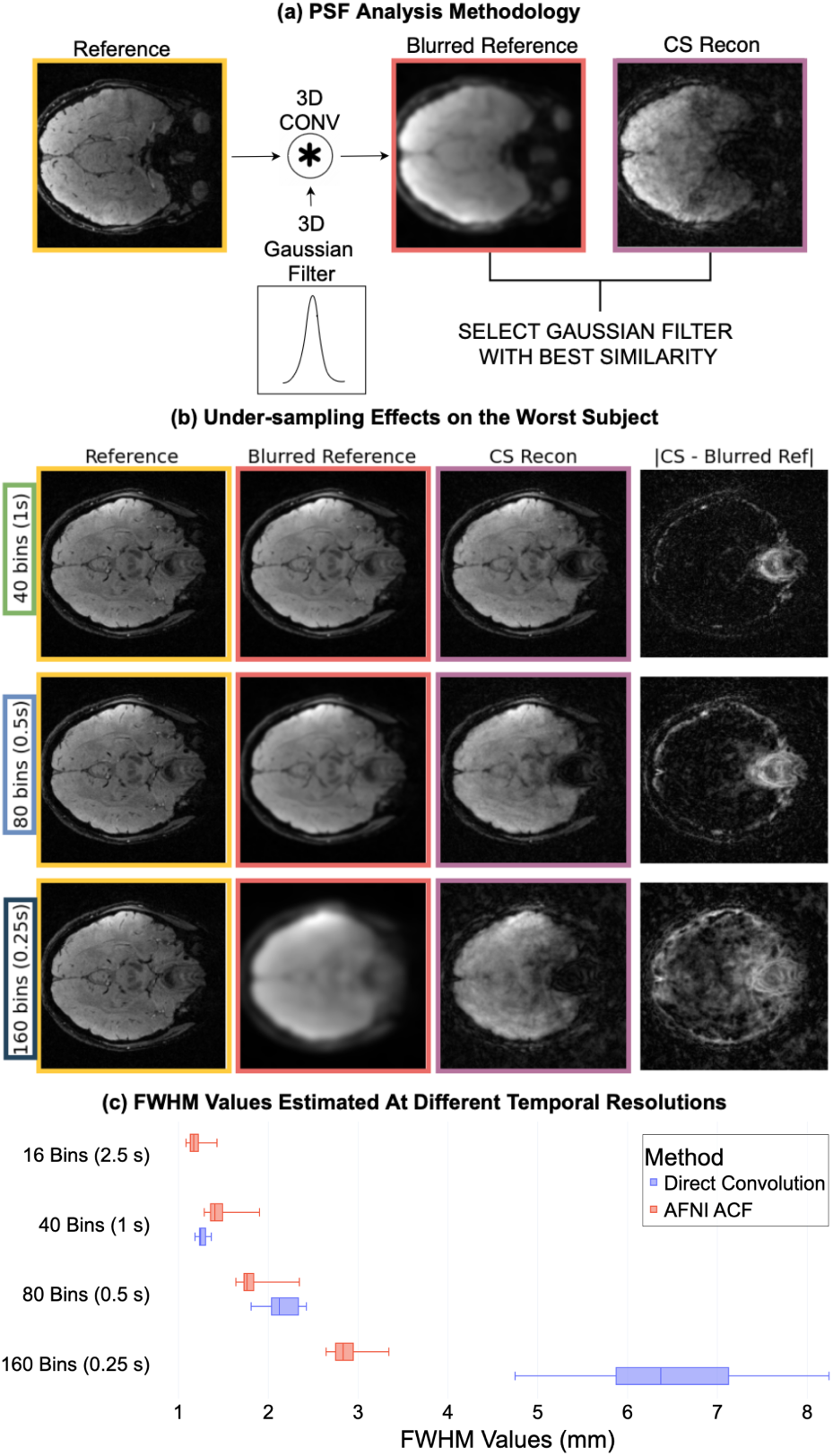
Point Spread Function (PSF) Analyses. **A)** Methodology Direct Convolution: We estimated the PSF using parametric fitting of a 3D Gaussian filter, optimizing the filter full width at half maximum (FWHM) to minimize the difference between the undersampled image (CS reconstruction with more than 16 temporal bins) and a reference image convoluted with the Gaussian filter. The 16-bin images were used as reference images. Each undersampled image was paired with the temporally closest reference image. **B)** Subject-Level Analysis: This section presents the reference image, the blurred reference generated using a Gaussian filter with the estimated FWHM, the undersampled image, and the absolute value of the difference between the blurred reference and the undersampled image for the subject with the highest estimated Gaussian blurring FWHM. This analysis helps determine whether CS reconstruction can be approximated as a 3D Gaussian smoothing operation or whether it introduces localized deformations in specific anatomical regions. The difference map (last column) shows that discrepancies are primarily concentrated at the image edges, particularly for lower undersampling rates (40 and 80 bins). However, at 160 bins, the undersampling effects deviate from simple Gaussian smoothing, even in central brain regions, and the CS reconstruction appears visually sharper while still exhibiting a high estimated blurring FWHM. This suggests that the smoothing introduced by CS reconstruction does not strictly follow a Gaussian model. **C)** Group-Level Analyses: The results of the PSF estimation at the group level with both methods confirm that FWHM increases with higher undersampling rates. Blue histograms show the PSF estimation performed with the Direct Convolution method, whereas red histograms present the PSF estimation as performed via the spatial autocorrelation of the GLM residuals using AFNI (AFNI ACF). AFNI’s PSF estimates increased from 1.17 mm at 2.5 s to 2.83 mm at 250 ms. Both methods estimate a similar FWHM at 1s and 0.5s. Moreover, this increase is nonlinear, suggesting that the undersampling level at 250 ms is approaching the critical undersampling rate for the adopted reconstruction method.

To further assess the impact of under-sampling on compressed sensing (CS) reconstructions, we estimated the PSF using an alternative image-based approach that operates directly on images reconstructed at different temporal resolutions, which we refer to as the direct convolution method. In this analysis, the PSF was quantified as the Gaussian blurring filter that best explains the differences relative to a reference image (the one reconstructed at 2.5s temporal resolution). Fig. 3 (a,b) presents the reference image, the blurred reference generated using a Gaussian filter with the estimated standard deviation, the under-sampled image, and the absolute difference between the blurred reference and the under-sampled image for the subject yielding the highest estimated blurring. The blue histograms in fig. 3, section C, illustrate a consistent trend across subjects, with FWHM values increasing with higher under-sampling. Specifically, the median FWHM values were 1.24 mm, 2.42 mm, and 6.37 mm for 1s, 500ms, and 250ms temporal resolutions, respectively. Two key observations emerge: (1) For lower levels of under-sampling (1s and 500ms temporal resolutions), image distortions relative to the blurred reference are primarily concentrated at the edges, with low impact on the region of interest (ROI). Notably, the observed differences are primarily localized to regions adjacent to the nasal cavities, which are well known to be affected by strong *B*_0_ inhomogeneities. (2) The inherent smoothing introduced by combining under-sampling and CS reconstruction increases with higher temporal resolutions (250ms), corresponding to higher levels of under-sampling. At a temporal resolution of 250 ms, we observe substantial image deformations that cannot be explained by Gaussian blurring alone and occur even within brain tissue. These effects contribute to increased volatility in the estimated percent signal change at higher temporal resolutions, making the PSF estimation with this image-based Gaussian model unreliable at 250ms resolution. To conclude, results show that both methods estimate a similar FWHM at 1s of 1.3mm and 0.5s of 2.1mm. For 250ms resolution, AFNI estimation shows a FWHM of 2.8mm, while the direct convolution method unreliably estimates 6.4mm.

### 2.3 Contributions of compressed-sensing and sparsity in T^∗^-weighted BOLD contrast in the visual cortex

Inference at the group level was computed via a one-tailed paired-t-test (*p <* 0.001, extended threshold of 100 contiguous voxels, dof = 19) and shows canonical activation in the calcarine sulcus as well as the lateral geniculate nuclei (41; 42). This is displayed in fig. 4, first row (∼23k (a.u) in voxel intensity of BOLD map threshold-free cluster enhancement (TFCE)-corrected (43)) which is based on the 33×2 reconstruction, i.e. 5D reconstruction performed by using the whole acquisition of all 33 trials (hereafter referred to as “5D-all”), see also fig. 6 for exemplar single-participant data from the lateral geniculate nuclei (supplementary material, fig. 6). The first three dimensions of the 5D-all reconstruction reflect space (voxels), the fourth dimension reflects the number of trials, and the fifth dimension reflects the estimated hemodynamic response (ON and OFF). The same analysis (one-tailed paired-t-test (*p <* 0.001, TFCE corrected, extended threshold of 100 contiguous voxels) has been performed on a single subject, i.e. between volumes corresponding to the ON phase of each trial and those corresponding to the OFF one, see supplementary material, fig. 7.

**Figure 4.**
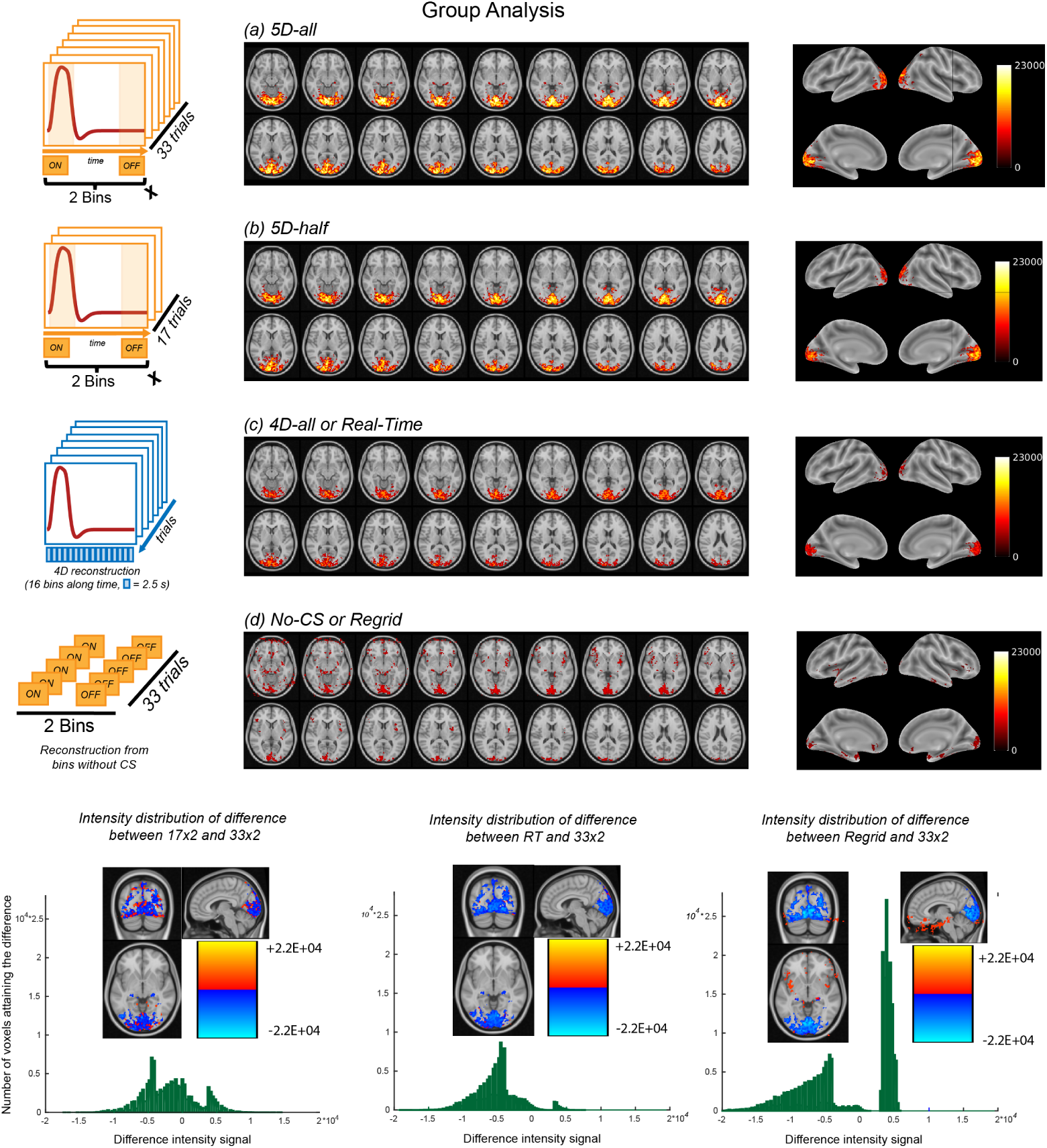
T_2_^∗^-weighted BOLD contrast and quantitative comparison of reconstruction approaches. Here, we present the inference at the group level computed as a one-tailed paired-t-test on images reconstructed with different numbers of trials (*p <* 0.001, extended threshold of 100 contiguous voxels). The first row presents results obtained by using all readouts (all 33 trials, 5D-all), where we can observe activation in the calcarine sulcus (∼23’000 (a.u) in voxel intensity of BOLD map TFCE-corrected, see colorbar). The second row contains the results for the 5D-half reconstruction, using half of the readouts, and the third row shows the 4D Real-time (4D-all) reconstruction (with all readouts). The maps show that we reach the same significance in the 5D-half case, as well as in the 4D-all, although with an observed decrease in the signal intensity. The fourth row presents the inference at the group level without the application of the compressed-sensing algorithm (No-CS). In this case, we observe highly non-uniform patterns (∼10’000 (a.u) in voxel intensity of BOLD map TFCE-corrected). The fifth row presents a histogram comparison where we plot in green the intensity of the difference between each of the performed reconstructions (5D-half, 4D-all, No-CS) relative to the reference reconstruction (5D-all). Blue voxels in the anatomical insets indicate either an activation present in the 5D-all reconstruction, but not visible in the other reconstructions, or voxels which activate in both reconstructions but at different intensities. Red voxels in the anatomical insets refer to activation present in the comparison reconstruction, but not in the 5D-all.

To further explore the role of under-sampling (the number of readouts used for reconstructing) and compressed-sensing (CS) (44), we report the group-level statistical inference also for the 17×2 reconstruction, a 5D reconstruction performed by using just half of the acquired trials (i.e., half of the readouts, fig. 4, second row; hereafter referred to as “5D-half”), as well as the group-level statistical inference for the T_2_^∗^-weighted BOLD contrast derived from the 4D real-time reconstruction (16 temporal bins, fig. 4, third row; hereafter referred to as “4D-all”, where the fourth dimension is the length of one trial chunked at different temporal resolutions), and finally from the reconstruction performed without the application of the CS algorithm (∼10k (a.u) in voxel intensity of BOLD map TFCE-corrected). In this case, the readouts are placed in a Cartesian grid, but no regularization is applied to explore shared information across nearby bins (fig. 4, fourth row; hereafter referred to as “No-CS”). In addition to the spatial pattern of significant activations, we also considered the computational time required across these different datasets. Table 1 reports the achieved reconstruction times, with up to 1h30 for the 4D-all.

**Table 1.** Reconstruction time of the different volumes. In this table we list the reconstruction times measured with a workstation equipped with two Intel Xeon CPUs, 512GB of RAM, and an NVIDIA Tesla GPU. All reconstructions were performed with *λ* = 0.01 and *ρ* = 0.06. *λ* is the regularization parameter of the objective function, whereas *ρ* is the convergence parameter of the alternating direction method of multipliers (ADMM). **CS** stands for Compressed-sensing, **TI** stands for Temporal Interval.

| Type of reconstruction | Time |
| --- | --- |
| 5D-all [3D volumes x **33 Trials** x 2 Activation (ON-OFF)] | 2h |
| no-CS [**no CS**: 3D volumes x **33 Trials** x 2 Activation (ON-OFF)] | 15min |
| 5D-half [3D volumes x **17 Trials** x 2 Activation (ON-OFF)] | 1h30m |
| 4D-all [3D volumes x Averaged TI: **2.5s** resolution] | 1h30m |
| 4D-all [3D volumes x Averaged TI: **1s** resolution] | 2h16m |
| 4D-all [3D volumes x Averaged TI: **0.5s** resolution] | 5h10m |
| 4D-all [3D volumes x Averaged TI: **0.250s** resolution] | 13h20m |

The spatial distribution of significant activations appears highly similar when using an identical statistical threshold (i.e. one-tailed paired-t-test, *p <* 0.001, extended threshold of 100 contiguous voxels) across the 5D-all, 5D-half, and 4D-all cases, although the activation maps show decreases in signal intensity. A very low and noisy signal is obtained when the compressed-sensing reconstruction is not performed (i.e., the No-CS reconstruction approach). These results, together with the PSF analyses, provide a preliminary indication of the role of under-sampling and CS reconstruction in reducing the amount of data that need to be used during reconstruction to retrieve comparable percent signal change within the acquired volumes.

To extend beyond these qualitative comparisons, we quantitatively compared, on a voxel-by-voxel basis, the spatial superposition and intensities of activations across reconstruction approaches with respect to the 5D-all method (see fig. 4, fifth row). Blue-toned voxels indicate where there was either an activation observed in the 5D-all reconstruction, which was not observed in the comparison reconstruction approach, or where activations were observed in both reconstructions, but with greater intensity in the 5D-all reconstruction. Blue voxels thus indicate where the 5D-all reconstruction outperformed the comparison reconstruction in extent and/or intensity. Red voxels, by contrast, indicate the opposite; i.e., where the comparison reconstruction outperformed the 5D-all approach. The histograms show these differences throughout the brain volume. The x-axis shows the difference in signal intensity, whereas the y-axis shows the number of voxels that attain such a difference.

Compared with the 5D-all reconstruction, the 5D-half approach displays activation patterns in the visual cortex, although with generally lower intensity (difference in voxel intensity between 0 and 10’000 (a.u) of BOLD map TFCE-corrected). Compared with the 5D-all reconstruction, the 4D-all approach also resulted in activation at generally the same loci, but with far lower intensity (fig. 4, center). Finally, comparing the 5D-all and No-CS approaches revealed that if CS is not applied, spurious patterns (red) appear throughout the volume, and the difference in the signal increases (25’000 (a.u) of BOLD map TFCE-corrected), see fig. 4, right.

## 3 Discussion

We introduced a new framework that aims at investigating BOLD fMRI from a different angle. We acquired whole-brain signals on a standard 3T clinical scanner while preserving a 1mm^3^ nominal spatial resolution, and reaching significant and reliable results at high (yet averaged) sub-second temporal resolution. Our approach allows organizing single k-space readouts in a flexible manner to retrieve specified signals at different temporal scales, which can be implemented after a single acquisition session, during reconstruction. We have shown that our acquisition and reconstruction strategies maintain the classical spatial resolution of traditional EPI-based approaches (15), while improving the recorded averaged temporal resolution at which brain activation can be studied, here up to 250ms, after scanning (see fig. 2 and 4).We performed two complementary PSF analyses to assess the effective spatial resolution across different temporal reconstructions: one based on AFNI’s spatial autocorrelation of GLM residuals, and another operating directly on the reconstructed images. Results show that both methods estimate a similar FWHM at 1s and 0.5s. For 0.25s, AFNI estimation shows a FWHM of 2.8mm. Importantly, the effective spatial resolution depends on both the chosen temporal resolution, but also on the number of repeated trials: the latter of which is controlled by the experimenter. Our 5D-all reconstruction approach was also able to detect significant activations at the group-level within thalamic structures (i.e. the lateral geniculate nuclei (41; 42; 45; 46), see fig. 6, while retaining whole-brain coverage, in addition to prominent clusters in the occipital lobe.

Up to now, measuring brain dynamics at a high temporal resolution with MRI (27; 47) was achieved only by compromising other factors (e.g., brain coverage in the case of laminar fMRI). For comparison, at a field strength of 3T as used in the present work, current multi-slice EPI approaches reach 0.5-1s with whole-brain coverage with 3mm^3^ spatial resolution (15), unless using accelerated methods that increase the risk and severity of artifacts. Our framework arguably mitigates the traditional trade-off between temporal and spatial resolutions by replacing it with a trade-off between the number of trials and spatial resolution, enabling the study of brain activation dynamics at novel high temporal resolutions and maintaining a full brain coverage. Further developments in the reconstruction algorithm adopting CS would be necessary to improve the temporal resolution below 250ms, while maintaining a reasonable amount of trials, as well as the spatial specificity of the signal.

From an analysis standpoint, our 4D reconstructions at different temporal resolutions do not require assumptions regarding the HRF shape, although prior knowledge on the stimulus onset across trials, in response to the visual stimulation, is used in the present study for the estimation of the T_2_^∗^ weighted activation maps. These activation maps can be computed either via a statistical analysis, or as a difference between the volume reconstructed by combining all readouts acquired during the activation state of the brain minus the volume reconstructed by combining all readouts acquired during the resting state of the brain. One pitfall of our methods is the current lack of integration of the GLM in our analysis pipeline, as we would need to inspect the single-trial temporal evolution, which is not currently possible in our framework. Further work to improve the CS reconstruction and radial acquisition scheme is therefore needed, in order to be able to expand the current applicability of the methods to more complex paradigms and event-related designs.

From the acquisition standpoint, spiral sampling patterns are less sensitive to motion and geometric distortions compared to Cartesian trajectories (EPI imaging), and have improved signal recovery, especially in frontal and parietal regions (11; 48). Here, we adopted a 3D radial - spiral phyllotaxis trajectory (35) for BOLD fMRI, and tested it as a component of our framework: we retrieve the desired activation pattern, validating our sequence’s use in BOLD functional imaging. While Radial-Cartesian schemes, as presented in (22; 49; 50) have been proved very successful in detecting BOLD fMRI responses at different magnetic fields, with an improvement in the overall image quality, our study currently lacks a systematic comparison with EPI schemes (only a single subject comparison is presented in fig. 6, top row). Further studies should explore these differences, as previously done in the literature for other Radial-Cartesian schemes. Furthermore, in such schemes, the flexible recombination of a variable number of consecutive (or not) acquired planes (or blades) does not allow for capturing averaged temporal dynamics at a fast pace. Temporal dynamics at the millisecond resolution have been observed via frameworks known as MREG (18; 19) or Inverse Imaging (20). Such techniques reach a temporal specificity approaching that observed with other neuroimaging modalities, such as MEG and EEG. The acquisitions are performed without (or very limited) gradient encoding and using the OVOC principle (one voxel one coil), exploiting the sensitivity of coils’ profiles and leveraging the repeated acquisition of identical k-space spokes. This results in a very high temporal resolution (100ms), but in a limited nominal spatial one. It would be very interesting to explore in future research novel acquisition techniques that could combine the spatial specificity of radial sequences with the temporal accuracy of MREG (or Inverse Imaging).

From the reconstruction standpoint, we validated the use of CS in retrieving the expected activation even when shortening the reconstruction (5D-half), reducing the acquisition time to 11 minutes, or when performing a quicker reconstruction (1h30m) to analyze the hemodynamic response function of the percent signal change (via the 4D-all reconstruction). Our results and those of other similar approaches in the field of CS applied to MRI (51; 52) suggest that the required number of trials (and therefore the computational time) could be further reduced in the future.

Another possible direction for future work is the application of this framework to situations in which flexible binning according to motion, trials, or physiological states may be informative. This could include studies involving participants who tend to move during scanning (e.g., patients, neonates, children, or elderly subjects), by allowing the exclusion of single readouts acquired during recorded motion. More generally, the approach may be applicable to imaging targets beyond the brain, for which MRI remains challenging due to breathing or cardiac motion (e.g., the spinal cord) (53; 54; 55), or in cases where motion is intrinsic to the system under study (56), such as the eyes or the neural layers of the retina, for which the shape of the hemodynamic response function (HRF) is not well characterized. This may be of particular interest when combined with techniques that explicitly resolve motion in these organs (57; 58; 59). With a sufficient number of trials, flexible binning of single k-space readouts may allow estimation of percent signal change in a new ROI without assuming a specific HRF shape, as shown in Fig. 2, since grouping is performed by trial prior to reconstruction. Once an HRF estimate is available, the data could then be re-binned to obtain T2^∗^ contrast maps.

Acquiring readouts at a high-sampling rate, might likewise help with investigating more deeply open questions regarding neuro-vascular coupling (60; 61; 62; 63; 64). In this way, BOLD activation will not only be coupled with physiology, but reconstructed according to physiological states, enabling the investigation of the contributors to hemodynamic activity. Another future direction for our framework could be the use of the same data for functional- and structural analyses (including cortical parcellations and surface inflation), by acquiring and consequently binning data according to different echoes. By alleviating the need to acquire separate structural and functional scans, this approach can avoid issues inherent to the co-registration of these different data types, which constitutes a potential source for error. This could be especially interesting for paradigms that require a large number of participants, and therefore a high number of individual data-quality checks (e.g., brain-wide association studies (65)). It may further allow creating regions-of-interest masks based on anatomical landmarks using the same data that is used for functional analyses, using independent criteria. It is our hope that, by introducing the Hi-Fi framework and outlining some of its potential applications, the broader research community will be encouraged to explore and adopt these strategies, thereby extending the usability of the framework.

## 4 Online Methods

### 4.1 Participants and visual stimulation

Twenty healthy volunteers (sex: 11 females, 9 males) underwent a single MRI session on a 3T clinical scanner (MAGNETOM Prisma*^Fit^*, Siemens Healthcare, Erlangen, Germany) equipped with a 64-channel head coil. Visual stimuli were back-projected onto a mirror attached to the head coil, with a total distance between the participants’ eyes and the screen of 102 cm. The study protocol received approval from the cantonal ethics committee (protocol number 2018 − 00240). The visual field on the mirror spanned 18°. The stimulation protocol followed a block design of 33 trials of 40s. This high number of trial repetitions for such a well-defined visual paradigm is justified by the fact that the technique is still in its prototype stage. Each trial entailed a 15s ON phase (active stimulation: 8Hz flickering checkerboard) followed by a 25s OFF phase (no stimulation: full-field and uniform grey image), see fig. 5, second row. This sequence allows the BOLD signal to reach its peak (after ∼5s) and to return to baseline (after ∼30s). A small centered attentional cue, changing color across time was placed at the center of the screen to help participants maintain fixation. Participants were asked to maintain the eyes relaxed and to minimize blinking and movement. Eye movements were tracked using an eye-tracking system (EyeLink 1000Plus, SR Research) synchronized with the MRI scanner via Syncbox (NordicNeuroLab). An Experiment builder (EyeLink) program was developed and used to control the calibration of the Eye-Tracker from outside the scanner room and to correctly synchronize the different hardware components of the experiment. Eye movement trajectories (of the right eye) were recorded using infrared light, with a sampling rate of 1000 Hz, through a mirror positioned inside the scanner bore, replacing the standard head-coil mirror usually available, which is not infrared compatible.

**Figure 5.**
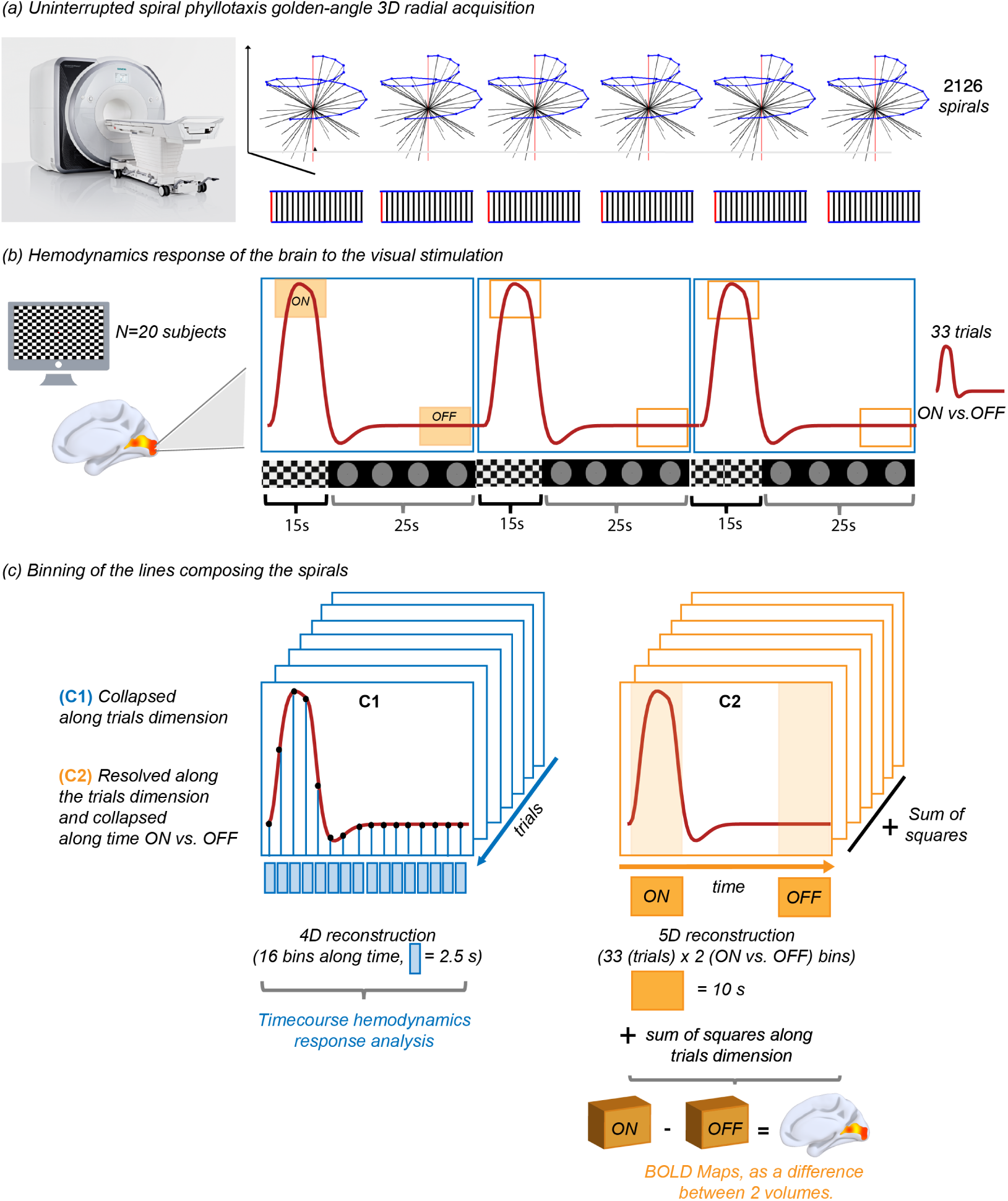
Overview of acquisition and reconstruction methods. **A)** The participant is placed in the scanner where we start acquiring data using a prototype uninterrupted gradient recalled echo (GRE) sequence with a 3D radial spiral phyllotaxis sampling trajectory, for a total of 46’772 readouts acquired. Each readout corresponds to either a red or black line in the plot below, with red lines indicating the first readout of each subspiral (commonly referred to as interleave). **B)** During the acquisition, the participant views a flickering checkerboard that follows a block design (i.e., 15s ON vs. 25s OFF). Roughly 5s after the visual stimulation onset, the hemodynamic response within the primary visual cortex reaches its peak. **C)** We performed different types of reconstructions. C1 - blue) refers to the temporal reconstructions: we averaged across all trials, every trial being divided into temporal bins of different temporal sizes: 2.5s, 1s, 500ms, 250ms. This produced a four-dimensional reconstruction (x, y, z, t), where the 4th dimension (t) refers to the averaged signal evolution curve. This allows to analyze the evolution of the percent signal change at different sub-second temporal resolutions. C2 - orange) refers to the 5D reconstruction: the ON-phase was reconstructed by retaining readouts acquired over the 5-15s post-stimulus time interval, while the OFF-phase by retaining readouts acquired over the 30-40s post-stimulus interval. All other readouts were discarded. *ON* and *OFF* 3D volumes across different trials were then unified via sum of squares, or retained to perform a single subject analysis, see 7. T_2_^∗^-weighted volume at the single participant level can be computed as the mathematical difference between the sum of squared *ON* and *OFF*, i.e. ||*ON* − *OFF*||.

### 4.2 Acquisition

Data were acquired using a prototype uninterrupted gradient recalled echo (GRE) sequence with a 3D radial - spiral phyllotaxis sampling trajectory (35; 66) that enables uniform k-space coverage in all bins, see fig. 5, first row. The acquisition and the visual stimulation were synchronized via a Syncbox (NordicNeuroLab). A total of 46,772 readouts (spokes=22, shots=2126) were acquired with a temporal resolution of 28.49 ms (i.e., the TR needed for a single k-space readout (spoke)), TE=25ms, FoV=192mm^3^ with 1mm^3^ isotropic nominal resolution (voxel size), FA=12°, and TA=22min. High spatial resolution anatomical T1 weighted data (MPRAGE, TE= 2.43ms, TR=1890ms, TI=955ms, FA=9°, FOV=256×256, 192 slices, voxel=1 mm^3^ isotropic) were acquired as the basis for ROIs segmentation.

### 4.3 T_2_^∗^-weighted BOLD contrast image reconstruction and signal extraction, 5D-all and 5D-half and No-CS

To assess the impact of different amounts of acquired readouts and the effect of regularization on the quality and reliability of the T_2_^∗^-weighted BOLD contrast images computed at the group level, we performed group-level statistical analyses on three different sets of reconstructions.

Three different 5D image reconstructions (32) were performed, where data were divided into five dimensions, x-y-z-rep-act, where rep refers to the number of trials (maximum 33) - the number of times the task was repeated, and act refers to the ON or OFF phase of the estimated hemodynamics response. The 5D image reconstructions were performed with different amounts of k-space readouts: the first reconstruction was performed using all acquired readouts (5D-all), the second reconstruction was performed using 50% of the acquired trials (5D-half), and finally a third reconstruction using all acquired readouts but without applying regularization (No-CS).

In all three 5D reconstructions, the ON-phase was reconstructed by combining all readouts acquired in 10-second intervals, starting 5 seconds after each stimulation, when the hemodynamic response is expected to reach its peak. In contrast, the OFF-phase was chosen to be temporally distant from any stimulation, using readouts acquired in a 10-second interval starting 30 seconds after the last stimulation, see fig. 5, block b (second row), orange boxes over the hemodynamic response curve (such ON vs. OFF volumes are reconstructed along the *act* dimension). All other readouts were discarded. The first two reconstructions were performed with a k-t-sparse SENSE algorithm(44) (image undersampling 20%), resulting in 66 3D volumes, see fig. 5, block c2, bottom right, orange reconstruction, first with the total number of readouts acquired and then with half of it. In such cases, total variation regularization (see table 1 for parameters and reconstruction time) was applied along the rep dimension (number of trials), as readouts acquired during the same hemodynamic phase, but across different trials, share information, while the different trials (N=33) were then combined via sum of squares. Rigid motion correction is achieved through a multistep process. First, temporally sequential ON and OFF volumes are reconstructed and used to estimate six rigid motion parameters per volume using SPM12 (67). This results in an estimation of 66 motion states. Motion estimates for each readout are approximated through linear interpolation. Then, the corresponding k-space corrections are applied to the raw k-space data using Fourier shift and rotation theorems. This process yields motion-corrected raw data, which are subsequently used as input for motion-corrected reconstructions. The T_2_^∗^-weighted volume at the single participant level can be computed as the mathematical difference between the sum of squared *ON* and *OFF*, i.e., ||*ON* − *OFF*||. Alternatively, see 7, a statistical inference can be performed at the single subject level between volumes corresponding to the ON phase of each trial and those corresponding to the OFF one (in this example, one-tailed paired-t-test (p < 0.001, TFCE corrected, extended threshold of 100 contiguous voxels). The inference at the group level was computed as a one-tailed paired-t-test (p<0.001, extended threshold of 100 contiguous voxels, dof = 19) between volumes corresponding to the ON phase of the participants and those corresponding to the OFF one. Such analyses have been performed in all three reconstructions mentioned above. We also computed histograms showing the distribution of intensity values of the differences between reconstructions. All reconstructions and analyses were performed in MATLAB (The MathWorks, Inc., Natick, MA, USA) on a workstation equipped with two Intel Xeon CPUs, 512GB of RAM, and an NVIDIA Tesla GPU. Time and parameters for the different reconstructions are presented in table 1.

### 4.4 Real-Time hemodynamic response image reconstruction, 4D-all

Temporal image reconstruction was performed to track the evolution of the percent signal change across the 40-second trial interval at different temporal resolutions. Therefore, every trial was subdivided into temporal bins of different temporal widths: 2.5s, 1s, 500ms, 250ms, thus producing a four-dimensional reconstruction (x, y, z, t) comprised of 16, 40, 80, 160 time frames (bins) respectively along the signal evolution curve. In single bins, data from different trials were grouped based on their relative timing with respect to the visual stimulation. After binning the data, the 3D volumes were reconstructed using a compressed-sensing-based algorithm (k-t sparse SENSE), where a Fourier transform operator was employed to exploit the shared information between different time frames (along the temporal dimension, see table 1 for parameters). The reconstructed images were used to extract the evolution of the hemodynamic response in regions of interest (ROIs) averaged across all trials, see fig. 5, block c1, bottom right, blue reconstruction. To retrieve the percent signal change at different temporal resolutions, a Fourier transform regularization was applied along the temporal dimension of the 4D-all reconstruction see fig. 5, block c1, bottom right, blue reconstruction. The hemodynamic response function has therefore been retrieved at different temporal resolutions, offline, after acquisition.

### 4.5 Percent signal change extraction and group-level analysis

Bilateral Regions of interest (ROIs) were extracted using FreeSurfer (68) at the individual participant level and included the calcarine cortex (encompassing the primary visual cortices), superior temporal sulci (encompassing the primary auditory cortices) and the postcentral gyrus (encompassing somatosensory cortices). Thalamic nuclei (including the lateral geniculate nuclei) were extracted from three exemplar individual participants (best percent signal change for these participants, see fig. 6). The average signal across different trials in the ROIs was computed for each of the temporal bins (16, 40, 80, 160 bins), after registering the T1 map from which the ROIs were derived onto our 4D-all reconstruction via SPM12 (67). We displayed the recorded hemodynamic response, see light purple curve of fig. 2, center, after outlier removal. After image inspection, we observed that two participants had corrupted reconstructed images, probably due to a misplacement of the head coils or heavy motion during scanning. Such participants were excluded only from the percent signal change analysis. Furthermore, the 4D 80 bins reconstruction of subject 20 resulted corrupted, so it could not be included in the PSF analyses. We performed two complementary PSF analyses to quantify the impact of undersampling and regularization on CS 4D-all reconstructions. As we did not include a retinotopic mapping sequence in our experimental protocol (69), we estimated the PSF at a post-processing level. One PSF analysis was based on AFNI’s spatial autocorrelation of GLM residuals, and another operating directly on the reconstructed images. The spatial autocorrelation of GLM residuals provides an absolute metric and it is proportional to the PSF autocorrelation (Wiener-Khinchin theorem, under the assumption that if the residuals are spatial uncorrelated, their autocorrelation is a Dirac delta function). It is possible to show that the measured autocorrelation of residuals provides an empirical estimate of the PSF’s effective spatial width, assuming the underlying residual field is spatially white and a specific model for the PSF shape. Functional MRI data were preprocessed using AFNI, including despiking and brain masking. Regression coefficients were estimated using REML, which accounts for temporal autocorrelations in the fMRI time series. Motion and physiological regressors, as well as baseline drift terms, were not included, since the reconstructions were already motion-corrected and the images were averaged across trials. Spatial smoothness of the residuals was then quantified using AFNI’s 3dFWHMx with the autocorrelation function (ACF) method. We reported FWMH of these estimated PSF kernels. In the second method, we implemented a parametric fitting of a 3D Gaussian filter. A range of Gaussian filters with varying widths was applied to generate blurred version of the reference images. The volumes generated with 16 temporal bins were considered as fully sampled and used as reference images. Then, we searched the optimal Gaussian filter width by minimizing the difference between the undersampled images (40, 80, and 160 bins) and the corresponding reference images blurred with the Gaussian filter. For each undersampled image, the width of the blurring filter was then estimated. The filter width that maximized the similarity between the blurred reference and the CS reconstruction was identified as the estimated PSF width, and retained as representative of the blurring effect. To pair reference images with the undersampled images, consider an example with 16 reference images and 160 undersampled images, where each reference image is paired with 10 consecutive undersampled images.

Additionally, to evaluate whether the inherent smoothing introduced by CS reconstruction can be well approximated by a Gaussian blurring kernel, we computed the absolute value of the difference between the blurred reference images and the CS reconstructions. Small differences represent a good approximation of the CS smoothing by a Gaussian blurring kernel. We reported FWMH of these estimated kernels.

## Acknowledgements

Financial support for this work has been provided by the Fondation Asile des aveugles (grant #232933 to M.M.M.), a grantor advised by Carigest SA (#232920 to M.M.M.), as well as the Swiss National Science Foundation (grants #220433 and #229214 to B.F., grants #169206 to M.M.M., #173129, #150828, and #143923 to M.S.). M.N. was supported by the Alexander von Humboldt Foundation and by the Intramural Research Program of the NIMH (ZIAMH002909). We thank Antoine Lutti and Quentin Reynaud for the fruitful discussions around the results of this article and its future applications.

## Authors contributions

B.F. conceptualized the problem with advice from M.M.M and E.F. B.F. and E.F. developed, implemented, and tested the protocol. B.F. performed the data analysis and image reconstruction with advice from M.N., J.J. and M.M.M. S.R., M.L. and M.D. set up the framework for image reconstruction. J.J. contributed to the introduction and state of the art. T.H., J.B. and T.K. provided technical support to derive the acquisition parameters at the scanner console, as well as to determine the sequence and its specifications. M.L. and C.W.R. performed the motion correction. G.D. performed the LGN extraction and contributed to the group-level analysis. M.L., R.v.H, M.S. and J.Y. contributed to formalizing the use of the compressed-sensing framework. G.P. contributed to the design of the network analysis techniques and interpretation. B.F., M.N., M.M.M. and E.F. drafted the manuscript, and all authors contributed to internal review.

## Supplementary material

**Figure 6.**
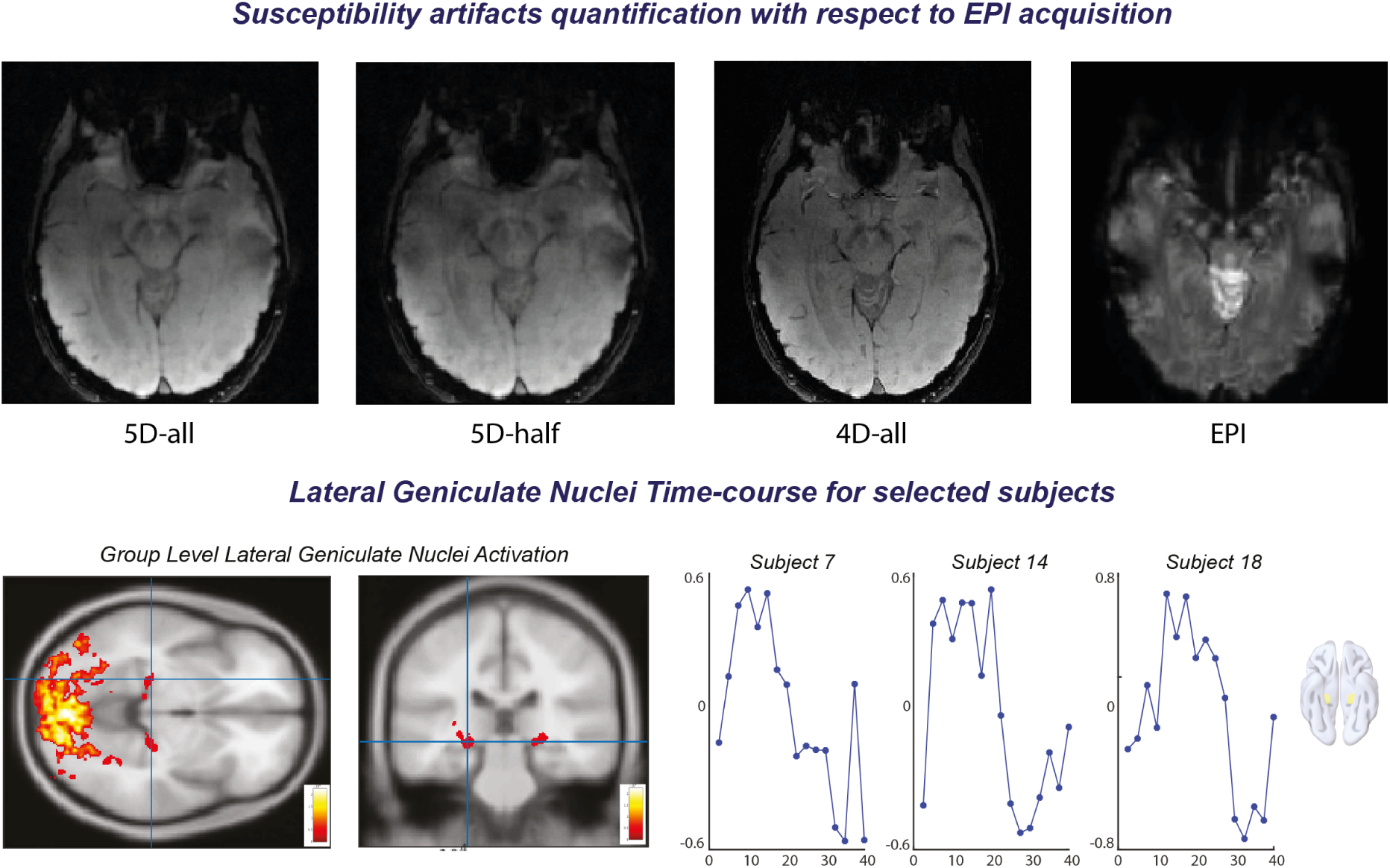
Top row: susceptibility artifacts quantification with respect to EPI acquisition. One subject underwent our protocol, together with an EPI (TR=2s, TE=73ms, FA=80°, resolution 2×2×3mm, TA=5.06min, Slices=14). A mask was created to isolate gray-matter in the selected axial slice after co-registration (the axial slice is therefore common to all four volumes, i.e. our three reconstruction and the slab acquired via EPI). We computed and report the coefficient of variation (CV) as a direct measure of the susceptibility artifacts magnitude across our reconstruction and the EPI. The measured CV is the following: 5D-all=0.407, 5D-half=0.435, 4D-all=0.351 and EPI=0.51. EPI reports the highest CV value. **Bottom row: activation in LGN.** In the 5D-all reconstruction we could observe hemodynamic responses in the lateral geniculate nuclei, i.e. thalamic subnuclei. We conducted an analysis at the single-subject level, reporting here the three participants with the strongest percent signal change in the LGN (around 0.8%).

**Figure 7.**
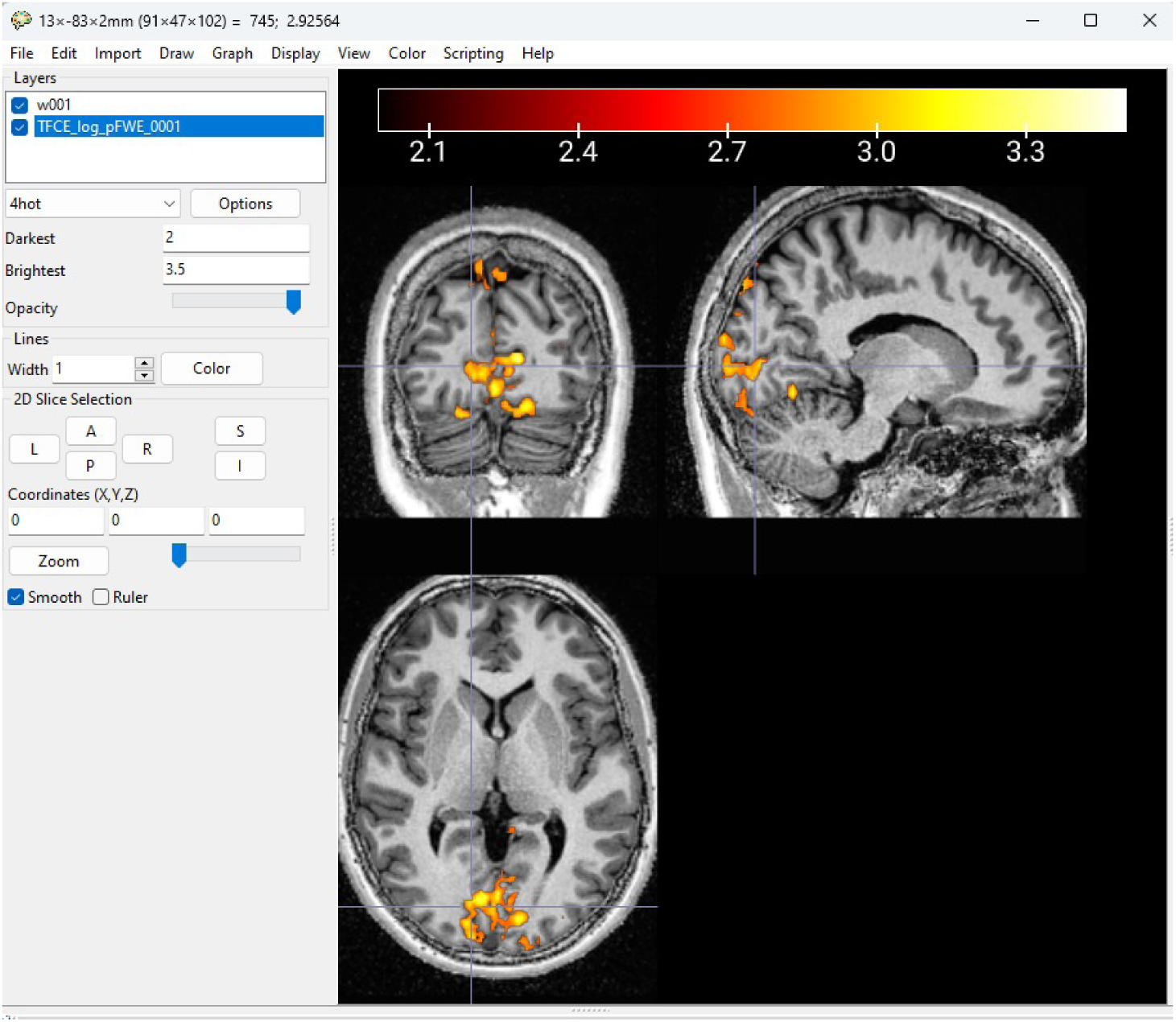
Single subject statistical analysis. Statistical inference can be performed at the single-subject level between volumes corresponding to the ON phase of each trial and those corresponding to the OFF one (in this example, one-tailed paired-t-test (p < 0.01 TFCE corrected, extended threshold of 30 contiguous voxels). This visualization is performed using MRIcroGL (70), where we show the interface and the thresholding of the log p equivalent map.

